# Inactivation of *Nphp2* in renal epithelial cells drives infantile nephronophthisis like phenotypes in mouse

**DOI:** 10.1101/2022.02.10.479981

**Authors:** Yuanyuan Li, Wenyan Xu, Svetlana Makova, Martina Brueckner, Zhaoxia Sun

**Author notes:** equal contribution authors (Yuanyuan Li, Wenyan Xu).

## Abstract

**Background:** Nephronophthisis (NPHP) is a ciliopathy characterized by renal fibrosis and cyst formation, and accounts for a significant portion of end stage renal disease in children and young adults. Currently no targeted therapy is available for this disease. *NPHP2* is one of the 25 NPHP genes identified to date. In mouse, global knockout of *Nphp2* leads to renal fibrosis and cysts. However, the precise contribution of different cell types and the relationship between epithelial cysts and interstitial fibrosis remains undefined.

**Methods:** Here, we generated cell-type specific knockout mouse models of *Nphp2* and characterized kidney morphology and phenotype. We additionally investigated the impact of removing cilia via deletion of the cilia biogenesis gene *Ift88* on phenotype severity in *Nphp2* mutants.

**Results:** Epithelial specific knockout of *Nphp2* in *Nphp2^flox/flox^; Ksp-Cre* mutant mice resulted in renal cyst formation and severe fibrosis, while *Nphp2^flox/flox^; Foxd1-Cre* mice, where *Nphp2* is deleted in stromal cells, displayed no observable phenotypes. Further, myofibroblast activation occurred early during disease progression and preceded detectable cyst formation in the *Nphp2^flox/flox^; Ksp-Cre* kidney. Moreover, concomitant removal of cilia partially suppressed the phenotypes of the *Nphp2^flox/flox^; Ksp-Cre* mutant kidney, supporting a significant role of cilia in *Nphp2* function *in vivo*.

**Conclusions:** Our results highlight the critical role of renal epithelial cilia and epithelial-stromal communication in NPHP.

**Significance statement:** Nephronophthisis (NPHP) is a ciliopathy characterized by interstitial fibrosis and epithelial cysts in the kidney. A significant portion of end stage renal disease occurring before age 30 results from NPHP. However, the molecular etiology of NPHP remains to be elucidated. By generating and analyzing tissue specific knockout mouse models of *Nphp2/Inversin*, authors pinpoint defective epithelial cells as the driver for both epithelial cysts and interstitial fibrosis. Moreover, the profibrotic response is triggered before cyst formation in the epithelial-specific *Nphp2* knockout model. Mechanistically, authors provide evidence that Nphp2 may function to inhibit a cilia-dependent pro-fibrotic and pro-cystic pathway.

## Introduction

Primary cilia are widely distributed cell surface organelles that function as signaling hubs for almost all vertebrate cell types^1–6^. Ciliary dysfunction is associated with an array of human disorders, collectively termed ciliopathies, that include obesity, mental retardation, retinal degeneration, and polycystic kidney disease (PKD), among many others (reviewed in^7^). In renal epithelial cells, cilia protrude into the lumen of tubules, exposed to both mechanical and chemical signals carried by extracellular fluid flow. Disruption of cilia biogenesis almost inevitably leads to the formation of epithelium-lined fluid filled kidney cysts in mouse^8, 9^. In addition, Polycystin 1 and 2 (Pc1 and Pc2), encoded by the autosomal dominant PKD (ADPKD) genes *Pkd1* and *Pkd2*, respectively, are targeted to cilia and this specific trafficking highly correlates with their *in vivo* function^9–13^. Interestingly, although both cilia biogenesis and Polycystin mutants develop cystic kidney, concomitant removal of cilia ameliorates cyst progression triggered by Polycystin inactivation, demonstrating that intact cilia are required for cyst growth in ADPKD models^14^. Based on these results, it was postulated that Polycystins function inside of cilia to inhibit a novel and uncharacterized cilia-dependent cyst activating (CDCA) signaling pathway^14^. This hypothesis implies the existence of a separate cyst-inhibitory pathway mediated by cilia, or a cilia-dependent cyst inhibiting (CDCI) pathway. In cilia biogenesis mutants, both CDCA and CDCI are removed, resulting in milder phenotypes compared to Polycystin mutants. Renal fibrosis is also frequently observed in cystic kidney diseases; but is often considered as secondary to cyst formation and the underlying mechanism remains poorly understood.

Nephronophthisis (NPHP) is a recessive renal cystic disease with cysts mostly confined in the corticomedullary junction region in the juvenile form of the disease, and more widely distributed across the kidney in the infantile form. Since multiple NPHP proteins are localized to cilia, basal bodies and/or show functional involvement in cilia biogenesis and/or signaling, NPHP is considered as a ciliopathy^15–22^. Compared with other types of renal ciliopathies, a prominent feature of NPHP is the severity of interstitial fibrosis. Genetically NPHP is highly heterogenous and at least 25 genes have been identified for the classic form of this disease (for a review, see^23^). However, mouse mutants of NPHP genes do not always show obvious renal phenotypes^22, 24, 25^. *Nphp2*, the homolog of which is associated with both infantile and juvenile NPHP in human, was discovered as a novel gene inactivated in a mouse mutant with reversed internal organ asymmetry along the left-right body axis and was originally named as *Inversin* (*Invs*)^26–28^. This whole-body knockout model additionally displays kidney cysts in both the medulla and cortex and renal fibrosis at the neonatal stage, resembling the renal phenotypes of infantile NPHP^28, 29^. However, the functional significance of *Nphp2* in different renal cell types is undefined. Moreover, the progression of renal fibrosis and its relationship with cyst formation remains to be characterized.

Here, to dissect tissue specific function of *Nphp2* and the mechanism of interstitial fibrosis resulting from defective *Nphp2*, we generated two conditional mouse knockout models: 1) targeting *Nphp2* deletion in epithelial cells of the distal nephron (*Nphp2^flox/flox^; Ksp-Cre* mice), and 2) deleting *Nphp2* in stromal cells *(Nphp2^flox/flox^; Foxd1-Cre* mice). We find that the NPHP-like phenotypes in mutant kidneys arise from abnormal signaling specifically within and from renal epithelial cells. Further, we investigated the relationship between cyst formation and interstitial fibrosis. Our results suggest that the fibrotic response precedes and therefore could initiate independently of cyst formation in this model. Mechanistically, we find that removal of cilia reduces the severity of *Nphp2* phenotypes, suggesting that, similar to Polycystins, Nphp2 inhibits a cilia-dependent cyst and fibrosis activating pathway.

## Results

### Inactivation of *Nphp2* in renal epithelial cells of the distal nephron leads to progressive cystic kidney disease in mouse

To dissect tissue specific function of *Nphp2*, we obtained embryonic stem cell clones generated by the European Conditional Mouse Mutagenesis Program (EUCOMM). The *Nphp2^tm1a^* “knockout-first” allele contains an IRES:lacZ-neo selection cassette located within two FLPE recombinase (FRT) sites and two loxP sites flanking the third exon of mouse *Nphp2* (Fig. 1A). Deletion between the two loxP sites is predicted to lead to frame shift and a premature stop codon. The *Nphp2^tm1a/+^* mice was crossed to a deleter strain expressing FLPe under the control of a ubiquitous actin promoter^30^ to generate a floxed allele of *Nphp2*. The carrier *Nphp2^flox/+^* mice were then crossed with carriers of *Ksp-Cre*, which directs Cre expression in the distal nephron of the developing metanephros after E11.5^31, 32^, to generate *Nphp2^flox/+^; Ksp-Cre mice*. Crossing *Nphp2^flox/flox^* with *Nphp2^flox/+^; Ksp-Cre* mice led to the generation of live *Nphp2^flox/flox^; Ksp-Cre* pups in Mendelian ratio. In mutants, neonatal pups showed progressive enlargement of the kidney: At postnatal day 7 (P7), the mutant kidney size was comparable to that in control animals and there was no significant difference in kidney to body weight (KBW) ratio between the two groups (Fig. 1B, C). At P14, the mutant kidneys were larger, and KBW ratios were significantly increased (Fig. 1B, C). Mutant kidneys became increasingly enlarged at P21 and P28 (Fig. 1B, C). We additionally compared KBW ratios between male and female mice and no significant differences were detected between genders in either control or mutant groups at P14 and P21 (Fig. 1D).

**Figure 1.**
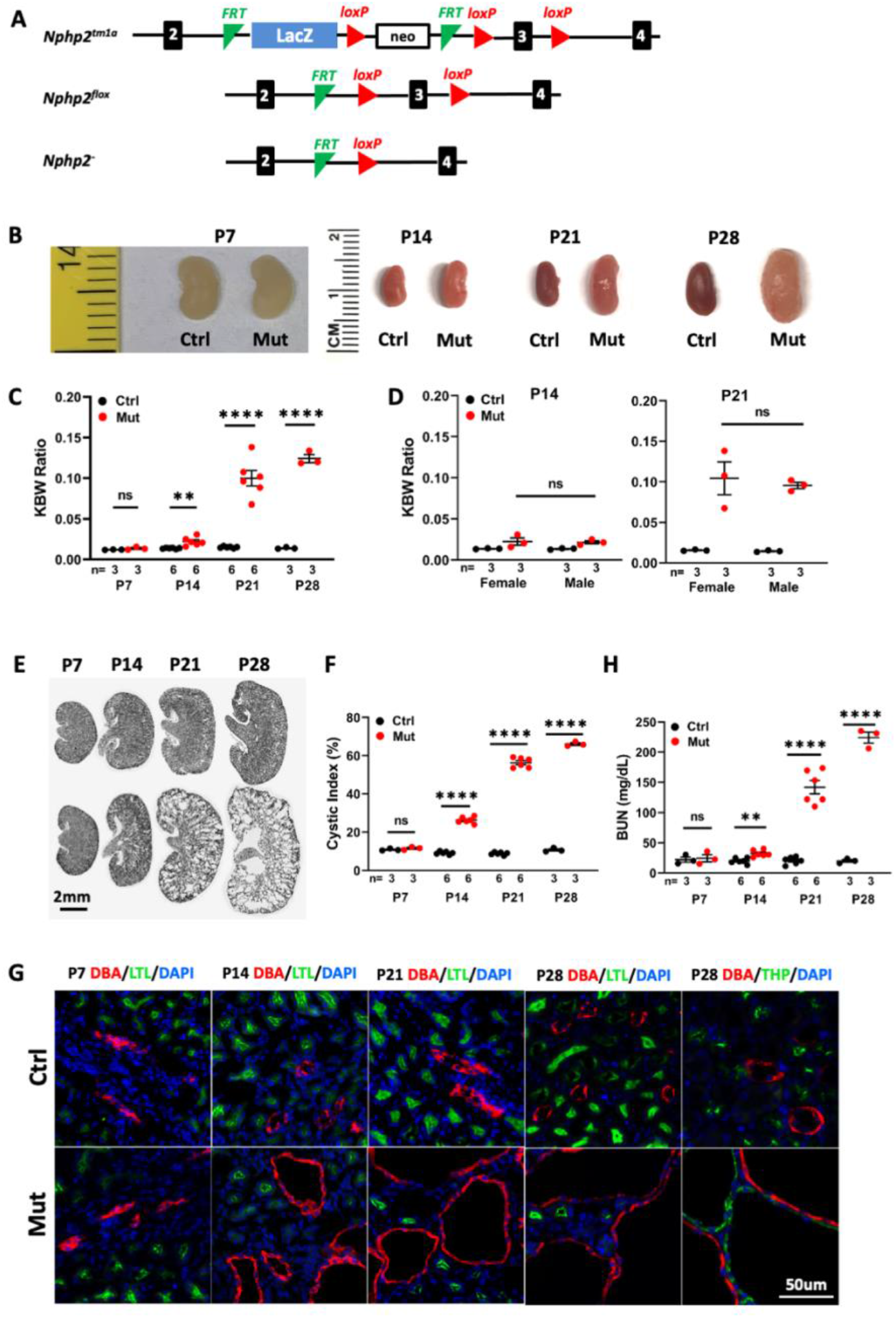
*Nphp2^flox/flox^; Ksp-Cre* mice develop progressive cystic kidney disease at the neonatal stage. (A) Generation of a floxed allele of *Nphp2*. FRT indicates FLPE recombinase sites, and loxP indicates Cre recombinase sites. (B) Gross morphology of the kidney from P7 to P28. (C) Comparison of KBW ratio in control and mutant mice. (D) Comparison of KBW ratio in male and female mice. (E) Hematoxylin/eosin-stained kidney sections from mutant and control mice. (F) Cyst formation quantified by cystic index. (G) Origin of cysts. DBA marks the collecting duct in red. The proximal tubule is labeled with LTL in green and the medullary thick ascending limb is labeled by THP in green. Nuclei are labeled with DAPI in blue. (H) BUN level in mutants and control mice. Ctrl: age matched control animals from the same cohort; Mut: *Nphp2^flox/flox^; Ksp-Cre* mutants; ns: not significant; **: p<0.01; ****: p<0.0001.

To inspect for potential cyst formation, we performed histological analysis on kidney sections and quantified cystic index as defined previously^33, 34^. Consistent with the progression of the KBW ratio, no cyst was detected in the P7 mutant kidney and the cystic index remained comparable between the wild type and mutant kidney (Fig. 1E, F). At P14, small cysts were detected, and the cystic index was already significantly increased in the mutant kidney (Fig. 1E, F). At P21, cysts became larger and more numerous in mutant kidneys (Fig. 1E), and by P28 mutant kidneys were severely cystic with large cysts throughout the cortex and medullar region (Fig. 1E). The cystic index progressively increased in mutant kidneys during the same time window (Fig. 1F).

To understand the cellular origin of kidney cysts in mutant mice, we performed immunohistochemistry, labeling different segments of the nephron in kidney sections. Consistent with histology, neither the collecting duct, labelled with Dolichos Biflorus Agglutinin (DBA), nor the proximal tubule, labelled with Lotus Tetragonolobus Lectin (LTL), showed enlargement at P7 (Fig. 1G). At P14, cysts were detected mostly in the DBA positive collecting duct (Fig. 1G). At P21, DBA positive cysts were enlarged and numerous. At P28, large cysts were detected in the thick ascending limb, labelled by anti-Tamm–Horsfall protein (THP), in addition to DBA positive cysts (Fig. 1G). By contrast, the LTL positive proximal tubule was free from cysts at P28 (Fig. 1G). Overall, the cystic region was consistent with the expression domain of *Ksp-Cre*.

To monitor kidney function, we assayed for the blood urea nitrogen (BUN) level. At P7, mutant and control mice were comparable (Fig. 1H). At P14, BUN levels became significantly higher in mutant mice (Fig. 1H), and differences in BUN levels became increasingly elevated at P21 and P28, demonstrating a progressive decline of kidney function (Fig. 1H).

Together, these results reveal a progressive renal cystic disease in *Nphp2^flox/flox^; Ksp-Cre* mice.

### *Nphp2^flox/flox^; Ksp-Cre* mice develop severe renal interstitial fibrosis

As interstitial fibrosis is a significant phenotype in NPHP, we first used trichrome staining to detect collagen deposition in P21 kidney sections. In mutant kidneys, an increase of the blue staining of collagen was detected (Fig. 2A). Further, we performed immunostaining of smooth muscle actin (SMA), a marker of myofibroblast, to monitor the fibrotic response over time. In the wildtype kidney, SMA signal was confined to the peri-vascular region. SMA signal was increased in the mutant kidney at P14 and became dramatically elevated and widespread in the mutant kidney at P21 and P28 (Fig. 2B).

**Figure 2.**
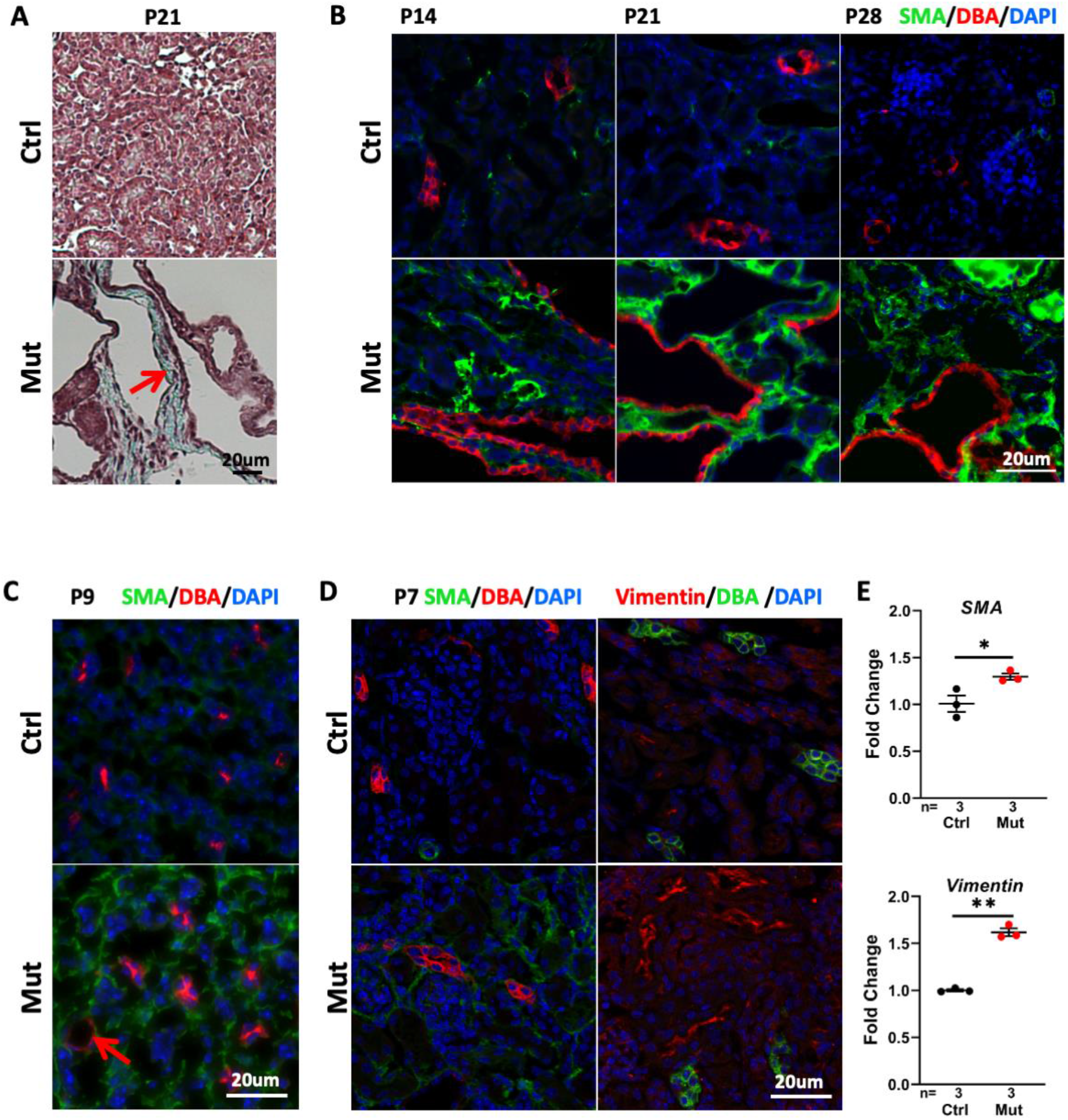
*Nphp2^flox/flox^; Ksp-Cre* mice develop early interstitial fibrosis in the kidney. (A) Trichome staining on kidney sections indicates blue collagen deposition (arrow) in a mutant at P21. (B) Increased signal of Sma in green in the cortex region of mutant kidneys from P14 to P28. DBA in red marks the collecting duct. DAPI in blue. (C) Increase of Sma staining (in green, DBA in red) at P9. DAPI in blue. Arrow points to a microcyst. (D) Modest increase of Sma (in green, DBA in red in the left panels) and Vimentin (in red in the right panels, DBA in green) in the cortex region of the mutant kidney at P7. DAPI in blue. (E) The level of *SMA* (upper) and *Vimentin* (lower) mRNA is increased in the mutant kidney assayed by Q-PCR using whole kidney lysates. Ctrl: age matched control animals from the same cohort; Mut: *Nphp2^flox/flox^; Ksp-Cre* mutants; *: p<0.05; **: p<0.01.

### Fibrotic response precedes cyst formation in *Nphp2^flox/flox^; Ksp-Cre* mice

To further delineate the relationship between cyst formation and interstitial fibrosis, we analyzed SMA signal at earlier stages. At P9, SMA signal in the cortex region was readily detected in mutant kidney sections, but isolated microscopic cysts were occasionally observed (Fig. 2C). At the non-cystic P7 stage, SMA signal was modestly increased in the cortex region (Fig. 2D). Quantitative PCR (QPCR) using whole kidney lysates also showed a significant increase of *SMA* mRNA level in the mutant (Fig. 2E).

To confirm the early activation of the fibrotic response, we examined the level of Vimentin, a marker of activated fibroblasts that becomes upregulated early during injury responses in the kidney^35, 36^. We first performed immunostaining using kidney sections. At P7, an increased signal was detected in the cortex region of the mutant kidney (Fig. 2D). QPCR using whole kidney lysates also showed a significant increase of *Vimentin* mRNA in the mutant (Fig. 2E).

Combined, these results reveal an increase of fibrotic markers at P7, preceding detectable cyst formation.

### *Nphp2^flox/flox^; Foxd1-Cre* mice show no observable renal phenotypes

As interstitial fibrosis is a prominent phenotype of NPHP, we investigated the role of *Nphp2* in the stroma by generating and characterizing *Nphp2^flox/flox^; Foxd1-Cre* mice. *Foxd1-Cre* drives Cre expression in metanephric mesenchymal cells destined to be stromal cells, including interstitial cells, mesangial cells and pericytes; and Cre activity can be readily detected at E11.5, similar to the onset of Ksp-Cre activity^31, 32, 37, 38^. The resultant mutant mice were viable. At P28, we did not observe any morphologic or functional phenotypes. No differences were seen in kidney size, KWB ratio, cyst formation, or collagen deposition in the mutant kidney at P28 (Fig. 3A-D, F). Moreover, immunostaining of SMA failed to detect any changes in the mutant kidney at P28 (Fig. 3G). In accordance, BUN levels were also normal in P28 mutants (Fig. 3E). Immunostaining of eGFP verified the wide expression of eGFP-Cre in the renal stroma in both the cortex and medulla region (Fig. 3H).

**Figure 3.**
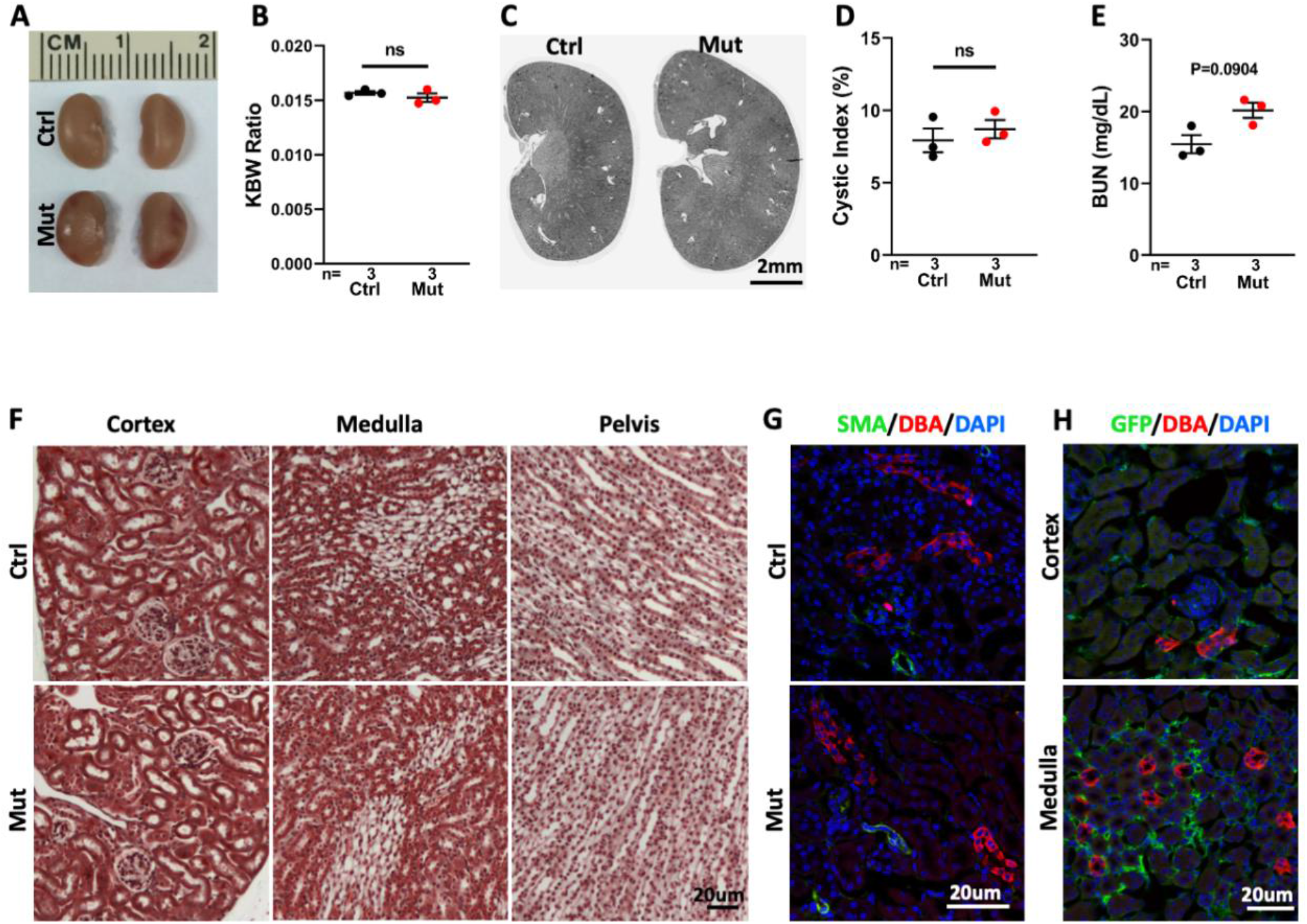
*Nphp2^flox/flox^; Foxd1-Cre* kidneys show no significant phenotype at P28. (A) Gross morphology of the kidney at P28. (B) KBW ratio in control and mutant mice. (C) Hematoxylin/eosin-stained sections of the kidney from mutant and control mice. (D) Cystic index in control and mutant kidneys. (E) BUN level in control and mutant mice. (F) Trichrome staining of kidney sections. (G) No increase of Sma (in green) can be detected in the mutant kidney. DBA in red labels the collecting duct. DAPI in blue labels the nucleus. (H) Immuno-staining of eGFP-Cre (green). DBA in red labels the collecting duct. DAPI in blue labels the nucleus. Ctrl: age matched control animals from the same cohort; Mut: *Nphp2^flox/flox^; Foxd1-Cre* mutants; ns: not significant.

### Genetic abrogation of cilia partially suppresses disease progression in *Nphp2^flox/flox^; Ksp-Cre* mice

Nphp2 is localized to and defines the Inversin compartment, a proximal segment of cilia^19,39^.However, it is also detected in subcellular locations outside of the cilium^40–42^. Moreover, Nphp2 is dispensable for cilia biogenesis^29, 43^. We validated this result in *Nphp2^flox/flox^; Ksp-Cre* mice. At P21, cilia were present in cyst-lining DBA positive cells as indicated by immuno-staining with the cilia marker anti-acetylated tubulin in the *Nphp2^flox/flox^; Ksp-Cre* kidney (Fig. 4A). In addition, Arl13b localization to the cilium was also undisrupted (Fig. 4B).

**Figure 4.**
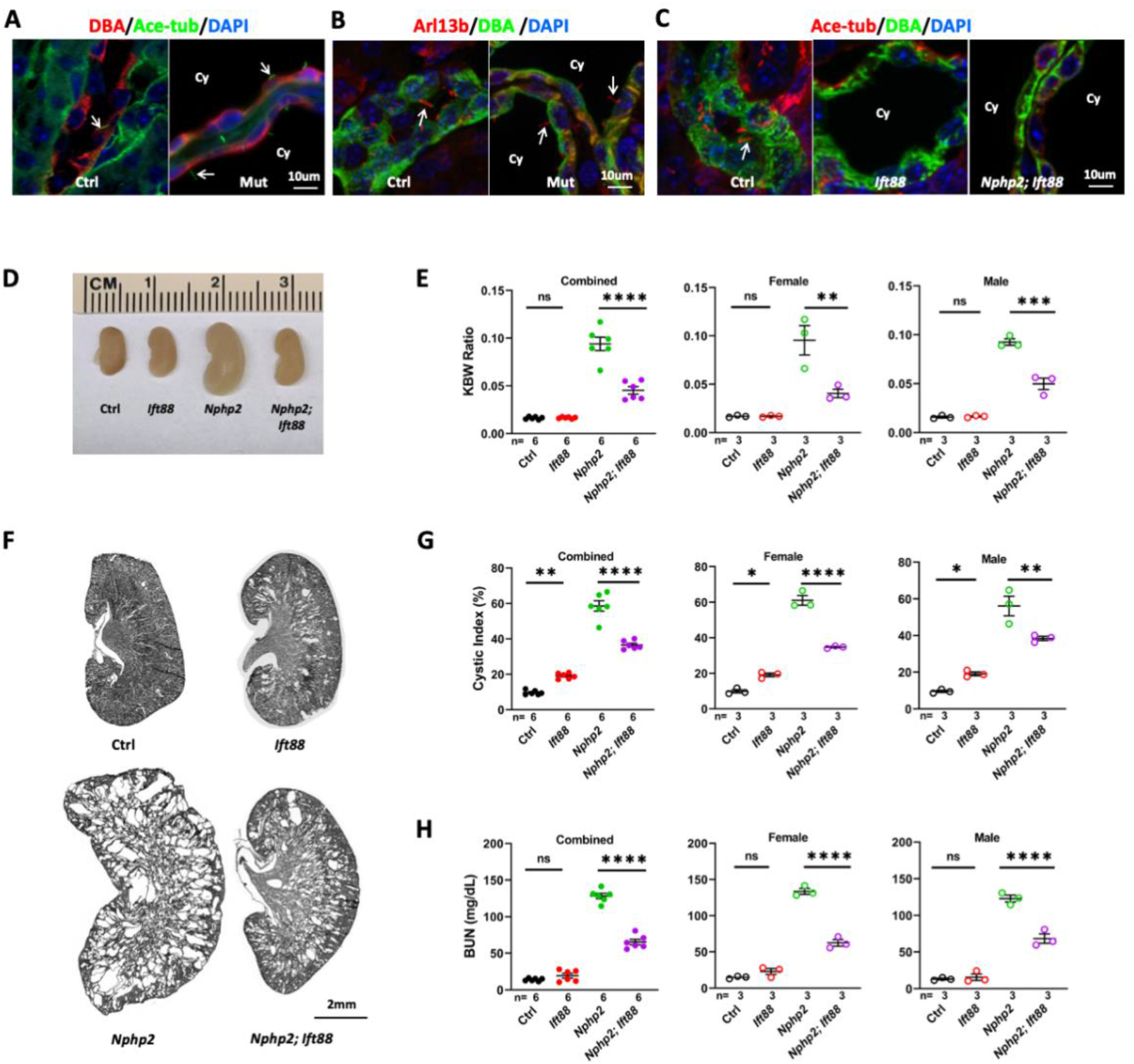
Genetic abrogation of cilia partially suppresses the phenotypes of *Nphp2^flox/flox^; Ksp-Cre* mice at P21. (A) *Nphp2* is dispensable for cilia biogenesis. Cilia (arrow) are labelled in green by anti-acetylated tubulin (Ace-tub). The collecting duct is labeled by DBA in red. DAPI in blue. (B) *Nphp2* is dispensable for ciliary localization of Arl13b labeled in red(arrow). DBA in green labels the collecting duct. DAPI in blue labels the nucleus. (C) Cilia biogenesis is disrupted in both *Ift88^flox/flox^; Ksp-Cre* single knockout and *Nphp2^flox/flox^; Ift88^flox/flox^; Ksp-Cre* double knockout mice. Cilia are labelled in green by anti-acetylated tubulin (arrow). The collecting duct is labeled by DBA in red. (D) Gross kidney morphology in control, *Ift88* single mutant, *Nphp2* single mutant and *Ift88; Nphp2* double mutant. (E) KWB ratio in control, *Ift88* single mutant, *Nphp2* single mutant and *Ift88; Nphp2* double mutant. (F) Hematoxylin/eosin-stained kidney sections of control, *Ift88* single mutant, *Nphp2* single mutant and *Ift88; Nphp2* double mutant. (H) Cystic index in control, *Ift88* single mutant, *Nphp2* single mutant and *Ift88; Nphp2* double mutant. (I) BUN level in control, *Ift88* single mutant, *Nphp2* single mutant and *Ift88; Nphp2* double mutant. Ctrl: aged matched control animals from the same cohort; *Nphp2: Nphp2^flox/flox^; Ksp-Cre* mutant; Ift88: *Ift88^flox/flox^; Ksp-Cre* mutant; *Ift88; Nphp2: Ift88^flox/flox^; Nphp2^flox/flox^; Ksp-Cre* double mutant; Cy: cyst. ns: not significant; *: p<0.05; **: p<0.01; ***: p<0.001; ****: p<0.0001.

The CDCA hypothesis posits that ectopic activation of a cilia-dependent cyst activating pathway underlies cyst formation in ADPKD^14^. If Nphp2 functions in a similar pathway, removal of cilia would rescue the phenotypes of *Nphp2* mutants. To test this hypothesis, we generated *Nphp2^flox/flox^;Ift88^flox/flox^; Ksp-Cre* double knockout mice. *Ift88* is an intraflagellar transport (IFT) gene essential for cilia biogenesis^8^. Immunostaining with the cilia marker anti-acetylated tubulin confirmed abrogation of cilia in the distal nephron in both *Ift88^flox/flox^; Ksp-Cre* single knockout and *Nphp2^flox/flox^; Ift88^flox/flox^; Ksp-Cre* double knockout mice, as expected (Fig. 4C). Consistent with previous reports, *Ift88^flox/flox^; Ksp-Cre* single knockout mice showed a slower progression of renal cystic disease in comparison to *Nphp2^flox/flox^; Ksp-Cre* mutants. At P21, neither kidney size nor KBW ratio were significantly changed in *Ift88^flox/flox^; Ksp-Cre* mutants compared to age matched control mice from the same cohort (Fig. 4D and E). In histological sections, *Ift88^flox/flox^; Ksp-Cre* mutant kidneys contained only small cysts but a significant increase of cystic index (Fig. 4F and G). By contrast, stage-matched *Nphp2^flox/flox^; Ksp-Cre* single mutants had enlarged and severely cystic kidneys, with significantly increased KBW ratios and cystic index (Fig. 4D-G). Interestingly, the increase of the KWB ratio and cystic index was reduced in *Nphp2^flox/flox^; Ift88^flox/flox^; Ksp-Cre* double mutant mice in comparison to *Nphp2^flox/flox^; Ksp-Cre* single mutant mice regardless of gender (Fig. 4E-G). Moreover, the BUN level was also significantly lower in *Nphp2^flox/flox^; Ift88^flox/flox^; Ksp-Cre* double mutants than in *Nphp2^flox/flox^; Ksp-Cre* single mutants regardless of gender, suggesting a partial suppression of kidney function decline (Fig. 4H).

Taken together, these results reveal that the phenotypes of *Nphp2^flox/flox^;Ksp-Cre* mutants is partially suppressed when cilia biogenesis is disrupted genetically.

## Discussion

Similar to infantile NPHP, mutants of the *ove* allele of *Nphp2/Invs*, generated by transgenic insertion of plasmid DNA in mouse, show severe cystic kidney disease and fibrosis at the neonatal stage^26–28^. It is possible that defective *Nphp2* in epithelial and stromal cells caused epithelial cysts and interstitial fibrosis, respectively. Alternatively, it is possible that abnormal epithelial-stromal cross-talks, resulting from tissue specific *Nphp2* inactivation, might have triggered the phenotypes observed in those previous studies.

In this study, we generated a floxed allele of *Nphp2* and used *Ksp-Cre* to specifically inactivate *Nphp2* in epithelial cells in the distal nephron and *Foxd1-Cre* to drive *Nphp2* deletion in the renal stroma. Significantly, inactivation of *Nphp2* in the distal nephron leads to cyst formation throughout the cortex and medullar region by P21, accompanied by interstitial fibrosis, recapitulating the main phenotypes of infantile NPHP. By contrast, inactivation of *Nphp2* in the renal stroma caused no observable phenotypes in the kidney at least up to P28, even though the onset of *Foxd1-Cre* expression in the developing metanephros occurs at the same time as that of *Ksp-Cre*^31, 32, 37, 38^. Combined, our results suggest that defective epithelial cells are the main driver of NPHP phenotypes in *Nphp2* mutants, highlighting the significance of epithelial-stromal cross-talks in the development and progression of interstitial fibrosis in this model.

PKD is frequently accompanied by renal fibrosis. In our previous study, we found that inactivation of the cilia biogenesis gene *Arl13b* in mouse leads to kidney cysts and fibrosis^44^. However, since the onset of fibrosis and cyst formation overlap in the *Arl13b^flox/flox^; Ksp-Cre* model, it is difficult to address whether interstitial fibrosis is solely secondary to epithelial cyst formation through, for example, mechano-stress exerted by expanding cysts. We therefore performed detailed time course analysis of cyst formation and the fibrotic response in the *Nphp2^flox/flox^; Ksp-Cre* kidney. Our results show that SMA and Vimentin is already upregulated at P7, before detectable cyst formation, suggesting that the fibrotic response is triggered before cyst formation by signals from defective epithelial cells in this model. Mechano-stress on stromal cells impinged by cysts, and additional signals from cystic epithelial cells could amplify the progression of fibrosis at later stages. Currently the nature of the signals from mutant epithelial cells remains unknown. Previous studies on the fibrotic response triggered by kidney injury detected upregulation of multiple developmental pathways, including TGFβ, Wnt, Notch and Shh (reviewed in^45^), providing ample candidate pathways for future analysis.

Previous work suggests that Nphp2/Inversin is specifically localized to a proximal segment of the cilium dubbed as the Inversin compartment^19^. Super-resolution imaging revealed that Nphp2 is localized to a novel fibril-like structure in the Inversin compartment^39^. Moreover, Nphp2 interacts with multiple infantile NPHP proteins, including Nphp3, Anks6 and Nek8; and it is required for concentrating these proteins to the Inversin compartment^19, 46^. On the other hand, Nphp2 is dispensable for cilia biogenesis and also found in the plasma membrane^29, 41^, mitotic spindle^40^ and stress granule^42^. It therefore remains possible that Nphp2 functions outside of the cilium. To test the relationship between cilia and Nphp2, we removed cilia genetically in the *Nphp2^flox/+^; Ksp-Cre* model by generating *Ift88^flox/flox^; Nphp2^flox/flox^; Ksp-Cre* double knockout mouse. Significantly, *Ift88* inactivation partially suppressed the severity of renal cyst and fibrosis, and improved kidney function in *Nphp2* mutants. This result suggests that, similar to Polycystins, Nphp2 may function to inhibit a cilia-dependent pro-cystic and pro-fibrotic pathway. Understanding the Inversin compartment could provide critical clues to the molecular mechanisms linking cilia, cyst formation and fibrosis. remains an interesting outstanding question.

## Concise Methods

### Animal Care Ethics

All mouse experiments were performed in Yale University School of Medicine in accordance with Yale University Institutional Animal Care and Use Committee guidelines.

### Mouse Breeding

Embryonic stem cell clones containing the *Invs^tm1a^* “knockout first” allele were purchased from EUCOMM and injected into C57/BL6J blastocysts to generate chimeric mice by Yale Genome Editing Center. Chimeric animals were mated to C57/BL6J mice to generate *Nphp2^tm1a/+^* mice. The *Nphp2^flox^* allele was generated by crossing *Nphp2^tm1a/+^* carrier mice with a deleter line expressing FLPe recombinase^30^. *Ksp-Cre* and *Pkhd1-Cre* mice^14, 32^ were kindly provided by the Somlo lab and were used to cross with *Nphp2^flox/+^* mice to generate *Nphp2^flox/+^; Ksp-Cre* or *Nphp2^flox/+^; Pkhd1-Cre* mice. *Nphp2^flox/flox^; Ksp-Cre* or *Nphp2^flox/flox^; Pkhd1-Cre* mice were obtained by crossing *Nphp2^flox/flox^* with *Nphp2^flox/+^; Ksp-Cre* mice or *Nphp2^folx/+^; Pkhd1-Cre* mice. *Foxd1-Cre* purchased from Jackson Laboratory (#012463) was used to obtain *Nphp2^flox/+^; Foxd1-Cre* and subsequently *Nphp2^flox/flox^; Foxd1-Cre* mice. The following primers were used for genotyping:

The floxed allele of *Nphp2*: 5’-AGGTTAGCAGCTGGGCAGGAT-3’ and 5’-TGAGGTAGA CAGTAGCATTCCTGC-3’;
The *floxed* allele of *Ift88*: 5’-GACCACCTTTTTAGCCTCCTG-3’ and 5’-AGGGAAGGGACTTAGGAATGA-3’
*Ksp-Cre*: For carrier: 5’-CAAATGTTGCTTGTCTGGTG-3’ and 5’-GTCAGTCGAGTGCACAGTTT-3’ Non-carrier: 5’-AGGCAAATTTTGGTGTACGG-3’ and 5’-GCAGATCTGGCTCTCCAAAG-3’
*Pkhd1-Cre*: 5’-CTGGTTGTCATTGGCCAGG-3’ and 5’-GCATCGACCGGTAATGCAGGC-3’
*Foxd1-Cre*, following protocol from Jackson Laboratory: 5’-TCTGGTCCAAGAATCCGAAG-3’, 5’-CTCCTCCGTGTCCTCGTC-3’ and *5’*-GGGAGGATTGGGAAGACAAT-3’.

### Immunostaining

Mouse kidneys were fixed by 4% PFA at 4°C overnight, embeded in OCT and cryo-sectioned at 5um. Sections were incubated in blocking buffer (10% FBS in PBST) for 1 hour at room temperature and then in primary antibody diluted with blocking overnight at 4°C. Slides were then washed three time with PBST for 10 minutes each, incubated in secondary antibody diluted with blocking buffer for 1-2 hours at room temperature protected from light, followed by washing three times with PBST, mounted in in ProLong Gold antifade reagent with DAPI (Molecular Probes by Life Technologies REF P36935) and imaged. The following primary antibodies and lectins were used: Rhodamine DBA (RL-1032; 1:100; Vector Laboratories), Fluorescein DBA (FL-1031; 1:100; Vector Laboratories), Fluorescein LTL (FL1321; 1:300; Vector Laboratories), THP (AF5175; 1:500; R&D Systems), Rabbit anti-Arl13b (17711-1-AP; 1:100; Proteintech), mouse monoclonal anti-acetylated tubulin (clone 6-11B-1; 1:5000; Sigma-Aldrich), mouse monoclonal anti-*α*-Smooth muscle actin (ab7817; 1:100; Abcam), Rabbit polyclonal anti-Vimentin (10366-1-AP; 1:100; Proteintech), mouse anti-GFP IgG (Roche, clone 7.1 and 13.1).

### Cystic Index Measurement

Cystic index was measured as previously published^34^. Briefly, two sections were analyzed for each animal. Hematoxylin/eosin-stained sections were scanned using the scan slide module in Metamorph v.7.1 acquisition software (Universal Imaging). Cystic and noncystic areas were measured by the Region Statistics feature in Metamorph. Cystic index is defined as cystic area to total kidney area in each section.

### Quantitative PCR Assay

For quantitative PCR (QPCR), mouse kidneys were lyzed in the Trizole reagent (Ambion by Life Technologies REF 15596018) and total RNA was extracted following the manufacturer’s instructions. Total RNA was used to synthesize cDNA using the SuperScript IV First-Strand Synthesis System (Invitrogen by Thermo Fisher Scientific REF 18091050). QPCR was performed using the KAPA SYBR FAST Universal (REF 07959389001) and BIO-RAD CFX96 Real-time System. The following primers were used:

Sma: 5’-CGTCCCAGACATCAGGGAGTA-3’ and 5’-ATAGCCACATACATGGCGGG-3’

Vimentin: 5’-ACCCTGCAGTCATTCAGACA-3’ and 5’-CAGAGAGGTCAGCAAACTTGGAC-3’.

GAPDH was used as internal control^47^.

### Statistical Analyses

t-test (and nonparametric tests) between paired samples (mutants vs control animals) and one-way ANOVA analysis between multiple groups were performed using GraphPad Prism 9.2.0.

## Acknowledgements

We thank members of the Brueckner laboratory, Somlo laboratory and Sun laboratory for helpful discussions; A. Cox for critical reading of the manuscript; S. Mentone for histology assistance; and D. Lonnette in George M. O’Brien Kidney Center at Yale for BUN analysis.

